# Predicting Future Forest Ranges Using Array-based Geospatial Semantic Modelling

**DOI:** 10.1101/009597

**Authors:** Elise Mulder Osenga

**Affiliations:** Science Writer, Earthzine

## Abstract

Studying the impacts of climate change requires looking at a multitude of variables across a broad range of sectors [1, 2]. Information on the variables involved is often unevenly available or offers different uncertainties [3, 4], and a lack of uniform terminology and methods further complicates the process of analysis, resulting in communication gaps when research enterprises span different sectors. For example, models designed by experts in one given discipline might assume conventions in language or oversimplify cross-disciplinary links in a way that is unfamiliar for scientists in another discipline.

Geospatial Semantic Array Programming (GeoSemAP) offers the potential to move toward overcoming these challenges by promoting a uniform approach to data collection and sharing [5]. The Joint Research Centre of the European Commission has been exploring the use of geospatial semantics through a module in the PESETA II project (Projection of economic impacts of climate change in sectors of the European Union based on bottom-up analysis).

---

The work of more than 40 scientists from 12 teams, the PESETA II project is intended to provide the European Commission (and other stakeholders) with an integrated analysis of the potential impacts of climate change in Europe [6]. This analysis takes the form of quantitative models that cover 10 main topics: agriculture, energy, river floods, droughts, forest fires, transport infrastructure, coasts, tourism, habitat suitability of forest tree species, and human health. The motivation behind these models is to assist adaptation plans for climate change impacts to Europe, while highlighting the uncertainties inherent in integrated assessment and in tipping-point phenomena [7, 8, 9].

**Figure 1:**
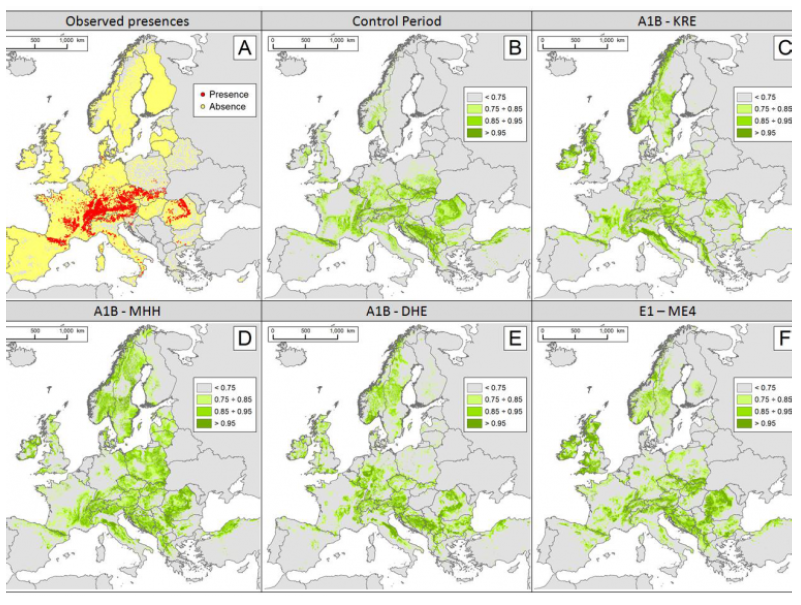
The Maximum Habitat Suitability based on the Relative Distance Similarity algorithm (RDS-MHS) estimated for Abies alba in the control period (B) and the climate change scenarios A1B (C,D,E) and E1 (F). Image Credit: Ciscar et al., 2014 [6], European Union, 2014.

The section on habitat suitability for tree species focuses on the European silver fir and can be used as an example of how the PESETA project works. The habitat suitability study incorporates data on existing tree distribution, bioclimatic factors, topography, and solar irradiation using a method known as the Relative Distance Similarity-Maximum Habitat Suitability Method (RDS-MHS).

The RDS technique calculates similarity between current geo-climactic conditions for a given area (such as Europe) and potential conditions for the same area likely under such climate scenarios as a 3.5 degree Celsius (A1B scenarios) or a 2 degree Celsius increase (E1 scenario).

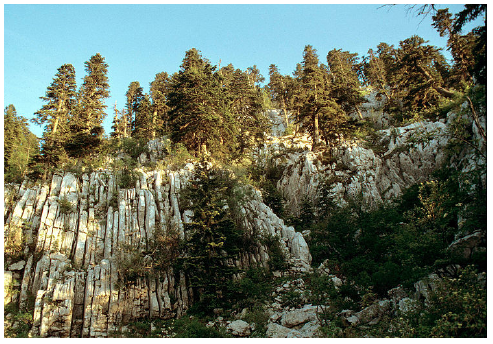
Dinaric calcareous Karst block fir forest in Orjen. Image Credit: Pavle Cikovac, Wikimedia Commons.

To create such model, the RDS software is programmed to define regions where silver fir currently lives or could live: the maximum habitat suitability (MHS) of the tree species. Next, the model uses this program to describe where the species could survive in the future, based on habitat predictions. The data are then used to create an index that represents the areas of new, lost, and stable habitat suitable under future climate conditions. These data could be applicable to planners by indicating areas where introduced silver fir would be most likely to survive.

The index also offers a projection of natural shifts in silver fir occurrence. Areas that have been calculated to have a high level of dissimilarity to current conditions may see a loss. By contrast, areas that are at a higher elevation or north of current populations of silver fir may see an increase or introduction of the species. Therefore, an index such as that of PESETA II could be valuable in helping land managers and others to determine which species are most likely to survive under future conditions and which may be futile to plant or restore.

The index is, however, limited in its efficacy as a management tool by two primary factors. First, the model assumes unlimited dispersal capacity by the trees under consideration. It does not account for actual limits to species migration. Second, since the models focus on climate response at a species, rather than an ecosystem level, complex habitat changes associated with changes in keystone species are not taken into account. For both of these reasons, the models tend to be conservative in their estimates of range loss.

In addition to its use for projecting future forest conditions, RDS has been applied to other environmental problems, from estimating soil erosion at the pan-European scale [10, 11] to deriving a fuzzy map of the Food and Agricultural Organization (FAO) ecological zones in Europe [12] and modelling susceptibility to landslides at the catchment scale [13].

Although the habitat suitability index and other RDS outputs are limited in their ability to forecast the future, they demonstrate a way of creating projects to help adaptation planning. The exact repercussions of climate change for local species and ecosystems remain uncertain, but by describing the maximum potential in future scenarios, models such as PESETA II can help planners to move forward.

